# Deciphering the Host-Pathogen Interface in COVID-19: The precision molecular insight into epitranscriptomic modifications of high-impact transcripts

**DOI:** 10.1101/2025.01.14.632961

**Authors:** Mateusz Maździarz, Katarzyna Krawczyk, Ewa Lepiarczyk, Łukasz Paukszto, Karol G. Makowczenko, Beata Moczulska, Piotr Iwanowicz, Piotr Kocbach, Jakub Sawicki, Leszek Gromadziński, Marta Majewska

## Abstract

The COVID-19 pandemic, caused by the severe acute respiratory syndrome coronavirus 2 (SARS-CoV-2), has had a profound global impact since its emergence in late 2019. Characterized by a wide spectrum of clinical manifestations, ranging from asymptomatic infection to severe respiratory distress and death, COVID-19 has necessitated extensive research into the host-pathogen interactions that drive disease progression. Understanding the molecular mechanisms underlying the host response to SARS-CoV-2 infection is crucial for the development of effective therapeutic interventions and preventative strategies. This study employed a multi-omic approach that combined direct RNA sequencing (DRS) and Illumina cDNA sequencing to investigate whole blood transcriptomic profiles in COVID-19 patients. By leveraging the unique capabilities of Nanopore DRS, which provides long-read sequencing data, we were able to capture not only gene expression levels but also crucial RNA modifications, including poly(A) tail length, non-adenine residue (non-A), pseudouridylation (psU), and 5-methylcytosine (m5C) methylation. This comprehensive analysis allowed us to identify differentially expressed genes (DEGs) and explore the impact of these RNA modifications on gene expression and function within the context of COVID-19. Our findings reveal significant alterations in gene expression patterns, poly(A) tail lengths, non-A and the prevalence of psU and m5C modifications in COVID-19 patients compared to healthy controls. These results provide valuable insights into the complex interplay between viral infection, host immune response, and RNA processing, contributing to a deeper understanding of COVID-19 pathogenesis.

## Introduction

The first documented case of COVID-19 emerged in Wuhan, China, in December 2019. Since then, the global case burden has surpassed 775 million, with over 7 million reported deaths (https://covid19.who.int/), which had unprecedented social and economic consequences. Advancements in prophylactic vaccination strategies have significantly mitigated the pandemic’s severity^1^, leading the World Health Organization (WHO) to declare the end of the COVID-19 public health emergency in May 2023 (https://news.un.org/en/story/2023/05/1136367). However, the emergence of novel variants with the potential to trigger surges in cases and mortality remains a concern, especially since the etiological agent responsible for COVID-19 disease, severe acute respiratory syndrome coronavirus 2 (SARS-CoV-2) ^2^, unlike other respiratory viruses, does not follow normal seasonal fluctuations, and waves of infection can happen at any time of year^3,4^. Its genome encodes a repertoire of viral proteins, categorized into non-structural proteins crucial for viral replication and pathogenesis and structural proteins essential for virion assembly^5,6^.

Despite the rapid global characterization of clinical symptoms associated with SARS-CoV-2 infection, a comprehensive understanding of the underlying host response and pathogenic mechanisms that govern disease progression toward recovery or fatality remains elusive. Elucidating the molecular foundations of COVID-19 pathogenesis is crucial for developing efficacious preventive and therapeutic strategies, ultimately aiming to reduce mortality and viral transmission. In our previous study, we investigated key genes engaged in SARS-CoV-2 infection using Illumina TruSeq RNA sequencing (RNA-seq) of peripheral blood samples collected from healthy donors and COVID-19 patients^78^. This time, to gain the most extensive insight into the whole blood transcriptomic profiles of the SARS-CoV-2-infected patients, we decided to exploit both RNA-seq short reads and Nanopore long reads. Nanopore sequencing technology enables direct, real-time analysis of long fragments of RNA in fully scalable formats. The transcripts obtained as a result of RNA long reads provide a range of valuable information, which is lost during the RNA-seq technique based on amplification, including RNA methylation, poly(A) tail sequence, and obtaining full-length transcript sequence^8,9^. This in-depth understanding of molecular changes induced by the virus is of utmost scientific and clinical importance, as it sheds light on the intricate interplay between viral infection, endothelial dysfunction, and immune responses in COVID-19. Such insights can guide the development of targeted therapeutic strategies to tackle the disease effectively.

## Results

### Chest computed tomography

The most frequently observed CT findings demonstrated a strong correlation with a COVID-19 diagnosis. These included ground-glass opacities, crazy paving with superimposed interlobular and intralobular septal thickening, consolidations, bronchovascular thickening in lesions, and traction bronchiectasis. The ground-glass opacities and consolidations are usually located bilaterally, they are diffuse and they show peripheral and basal distribution. Peripheral distribution of the lesions, ground-glass opacities, and bronchovascular thickening in the lesions were found to have the highest value in diagnosing COVID-19 patients (Fig. 1).

**Fig. 1.**
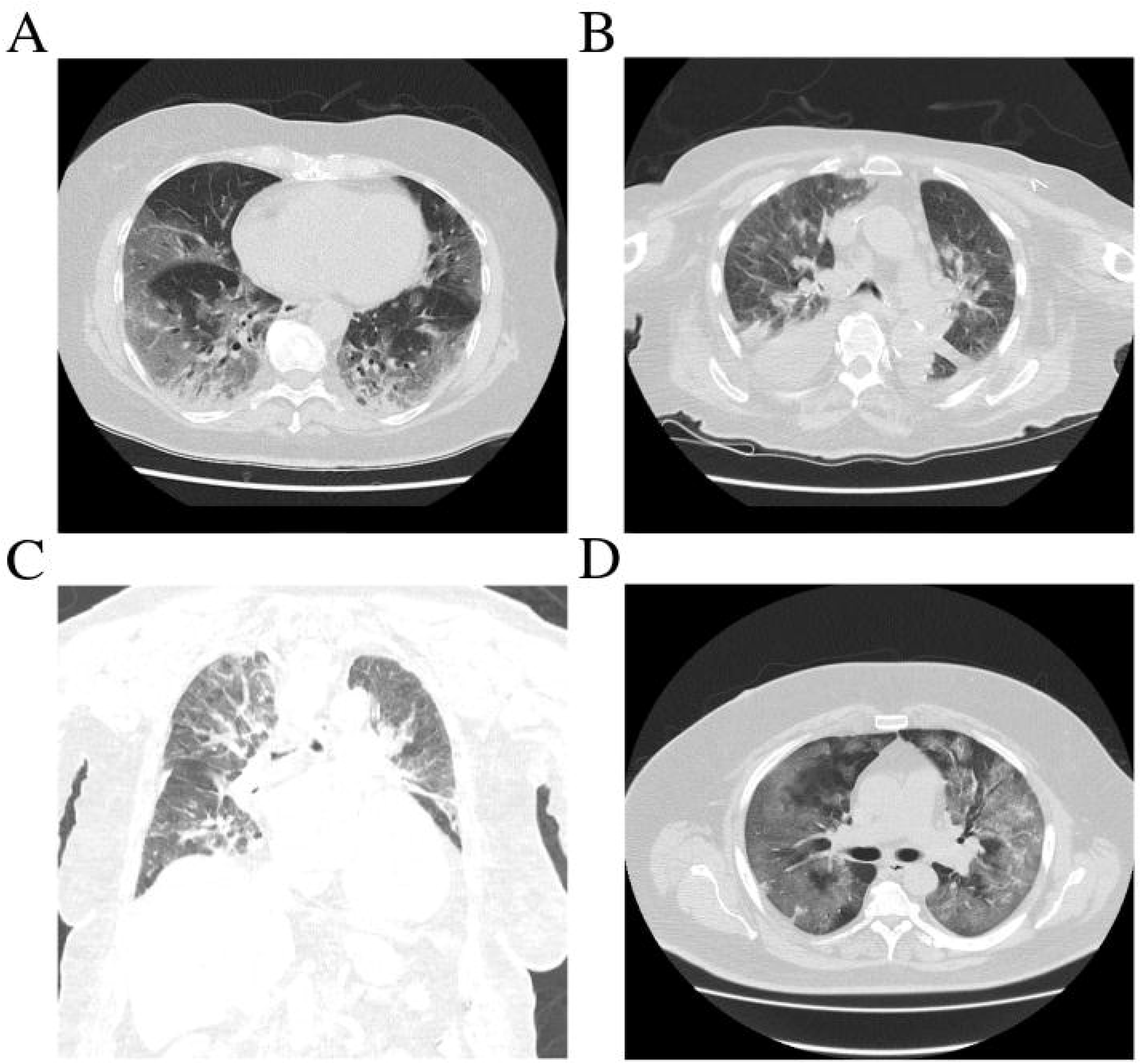
Noncontrast axial and coronal plane chest CT images of extensive inflammatory changes characteristic of COVID-19 disease. **A** CT images of a 67-year-old woman with Covid-19 for 7 days with mild COVID-19 pneumonia. Extensive ground-glass opacities with consolidation, primarily in the lower lobes, were noted, consistent with severe COVID-19 pneumonia. Approximately 50% of the lung parenchyma was affected. **B** CT images of a 90-year-old woman with Covid-19 for 2 days with mild COVID-19 pneumonia. Extensive bilateral ground-glass opacities with marked interlobular septal thickening, consistent with severe pulmonary involvement due to COVID-19 or pulmonary edema. Approximately 90% of the lung parenchyma was affected. Subsegmental atelectasis or consolidation was present in the lower lobes. Significant pleural effusions were noted, measuring up to 5 cm on the right and 2 cm on the left. Mediastinal lymphadenopathy was present, with nodes measuring up to 16 x 11 mm. **C**, **D** CT images of a 63-year-old man with Covid-19 for 7 days with mild COVID-19 pneumonia. The lungs exhibited marked hypoventilation with extensive, confluent ground-glass opacities and consolidations involving all lobes. Patchy, dense consolidations were most prominent in the upper lobe of the left lung, characteristic of severe COVID-19. Approximately 70-75% of the right lung parenchyma and 75-80% of the left lung parenchyma were affected.

### Comparative transcriptome analysis of COVID-19 patients reveals distinct gene expression signatures

Direct RNA sequencing (DRS) provided information on the expression of 16,510 genes, of which 197 were identified as differentially expressed genes (DEGs). Among these molecules, 112 were downregulated and 85 were upregulated. The log2FoldChange ranged from -7.78 to 4.23 for all molecules (Fig. 2A-D, Supplementary Table 1). Subsequently, a GO (Gene Ontology) analysis of all significant molecules was conducted, which were significantly involved in immune response (GO:0006955), response to virus (GO:0009615), and defense response (GO:0006952) (Supplementary Table 2). In the next stage of bioinformatics, cDNA expression generated by Illumina was analyzed. Sequencing using this method provided information on 36,689 genes. A total of 707 molecules were classified as DEGs, including 362 downregulated genes and 345 upregulated genes. The log2FoldChange ranged from -11.41 to 9.36 for all genes (Fig. 2A-D, Supplementary Table 3). Subsequently, a GO analysis of all significant molecules was conducted, which were significantly involved in immune response (GO:0006955), response to virus (GO:0009615), and defense response (GO:0006952) (Supplementary Table 4). The next step involved plotting the Pearson correlation between the log2FoldChange of common genes. A total of 52 DEGs common to the cDNA and DRS methods were classified. Among the common genes, *NFIX*, *LAP3*, *IGLC3*, *SAMD9L*, and *IFIT3* were identified (Fig. 2A,C). The correlation coefficient between their expression log2FC values was 0.96, with a correlation P-value of < 2.2e-16 and a 95% confidence interval of 0.9393051 to 0.9797827. It was found that similar expression signatures were indicated by the results of the correlation coefficients for common genes (Fig. 2B).

**Fig. 2.**
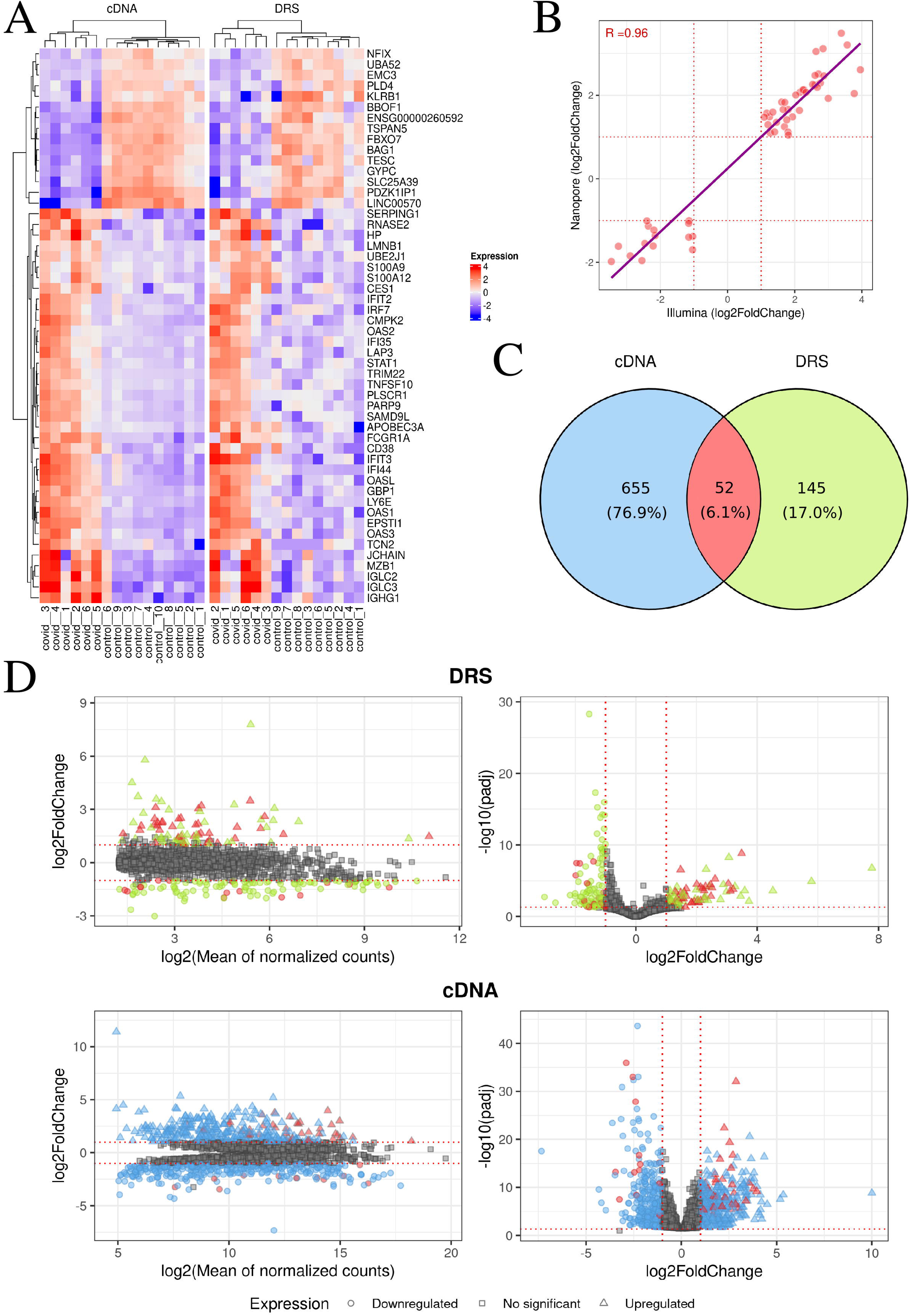
Gene expression profiling of controls and COVID patients. **A** The heatmap displays the normalized expression of 52 common genes for DRS and cDNA sequencing. Red color indicates values greater than 0, while blue indicates values less than 0. **B** The scatter plot illustrates the relationship between log2FoldChange values for the 52 DEGs identified in both sequencing methods. The x-axis represents log2FoldChange values for cDNA Illumina sequencing, while the y-axis depicts log2FoldChange values for DRS Nanopore sequencing. The red line highlights the Pearson correlation. The R value is displayed in the upper left corner. **C** The Venn diagram presents DEGs in cDNA Illumina (blue), DEGs in DRS Nanopore (green), and common DEGs (red). **D** DEGs characteristic of COVID-19 patients. The top panels correspond to DRS Nanopore, while the bottom panels represent cDNA Illumina sequencing. All DEGs are colored according to the Venn diagram. On the left, MA plots show the relationship between log2FoldChange and log2 (Mean of normalized counts), while on the right, Volcano plots show the relationship between -log10(padj) and log2FoldChange.

### Dissecting the mRNA landscape of COVID-19: A role for poly(A) tail dynamics

A total of 2,029,252 poly(A) tails were analyzed. A statistically significant difference in the global distribution of poly(A) tail lengths was found between COVID-19 and control patients (P-value < 2.2e-16). 6,524 transcripts with poly(A) tails were identified (Fig. 3A, Supplementary Table 5). Lengthening of poly(A) tails was observed in as many as 879 genes in COVID-19 patients, compared to only 8 in the control group (Fig. 3G). Subsequently, a GO analysis of all significant transcripts was conducted, which were significantly involved in immune response (GO:0006955), response to virus (GO:0009615), and defense response (GO:0006952) (Supplementary Table 6).

**Fig. 3.**
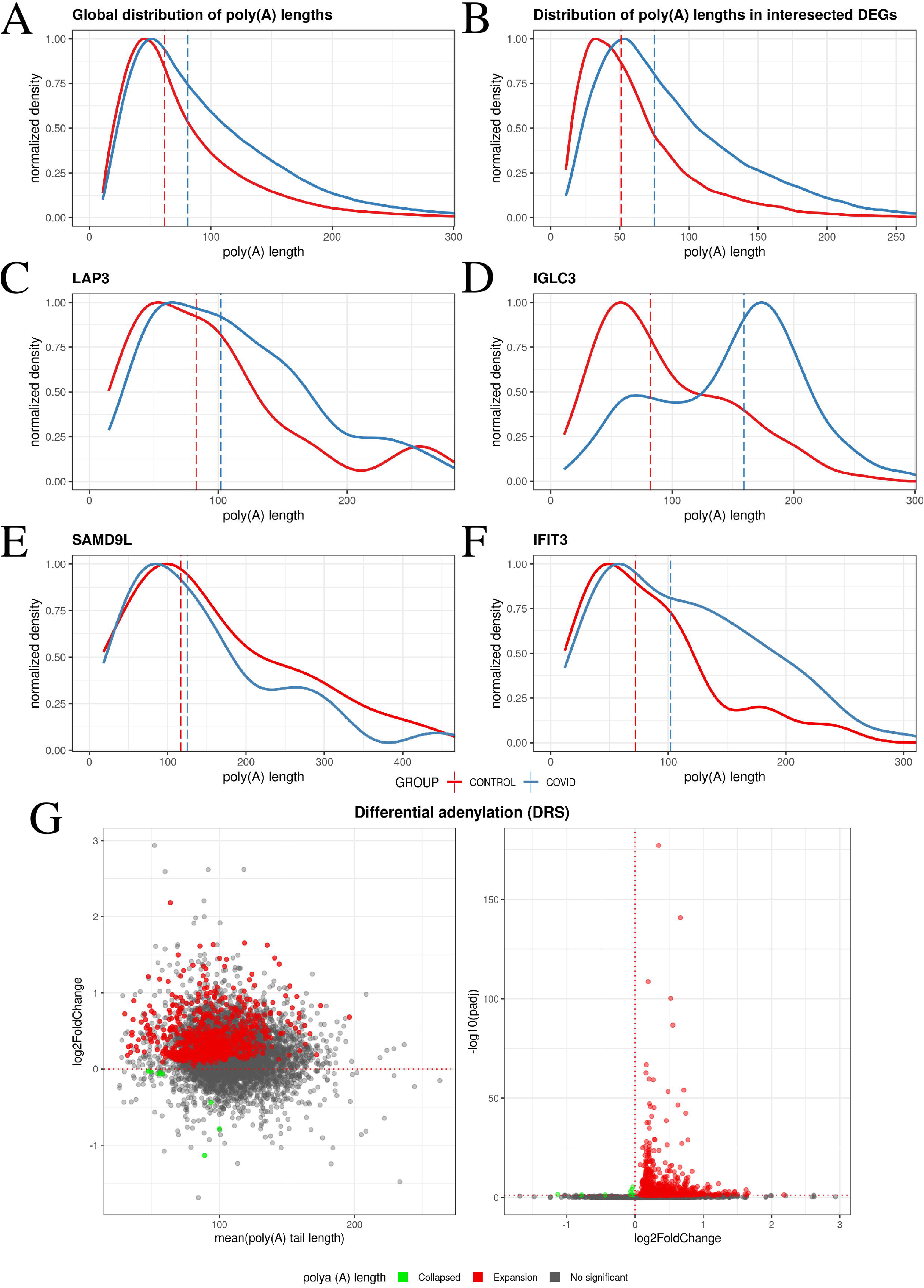
Poly(A) tail length variation across the genes. **A-F** The density plots depict the normalized transcript density on the y-axis and the length of poly(A) tails on the x-axis. Panel A shows the global distribution of poly(A) tails, panel B shows the poly(A) distribution of intersected DEGs, while panels C to F present the poly(A) distribution for specific genes: *LAP3*, *IGLC3*, *SAMD9L*, and *IFIT3*, respectively. **G** Illustrates genes with distinct poly(A) characteristics. The left panel displays a modified MA plot, depicting the relationship between log2FoldChange (based on poly(A) tail length) and the mean poly(A) tail length. Conversely, the right panel presents a Volcano plot, showcasing the relationship between log2FoldChange (based on poly(A) tail length) and - log10(padj). Significant genes are highlighted in blue, with shortened tails represented by circles and lengthened tails denoted by squares.

Furthermore, differences in poly(A) tail lengths in differentially expressed genes (DEGs) common to both methods were examined. A significant result was obtained, indicating a lengthening of poly(A) tails in DEGs of COVID-19 patients (P-value < 2.2e-16) (Fig. 3B). Additionally, differences in poly(A) tail lengths were examined for individual genes: *LAP3* (P-value = 0.046) (Fig. 3C), *IGLC3* (P-value < 2.2e-16) (Fig. 3D), *SAMD9L* (P-value = 0.7385) (Fig. 3E), and *IFIT3* (P-value = 0.026) (Fig. 3F).

In addition to adenine, other nucleotides such as guanine, uracil, and cytosine were identified within poly(A) tails. A global analysis of the frequency of non-adenine (non-A) residues was conducted, revealing that guanine (22,129 for COVID-19 patients and 15,447 for control patients) was found more frequently poly(A) tails than cytosine (20,300 for COVID-19 patients and 13,165 for control patients) or uracil (19,141 for COVID-19 patients and 12,576 for control patients) (Fig. 4A, Supplementary Table 7). A shift in this trend was observed in DEGs common to both methods. Cytosine (911) was determined to be the most frequent non-A residue in COVID-19 patients, whereas uracil (556) was found to be the most frequent in the control group. Guanine (700 for COVID-19 patients and 357 for control patients) was identified as the least frequent non-A residue in both groups (Fig. 4A,B, Supplementary Table 8).

**Fig. 4.**
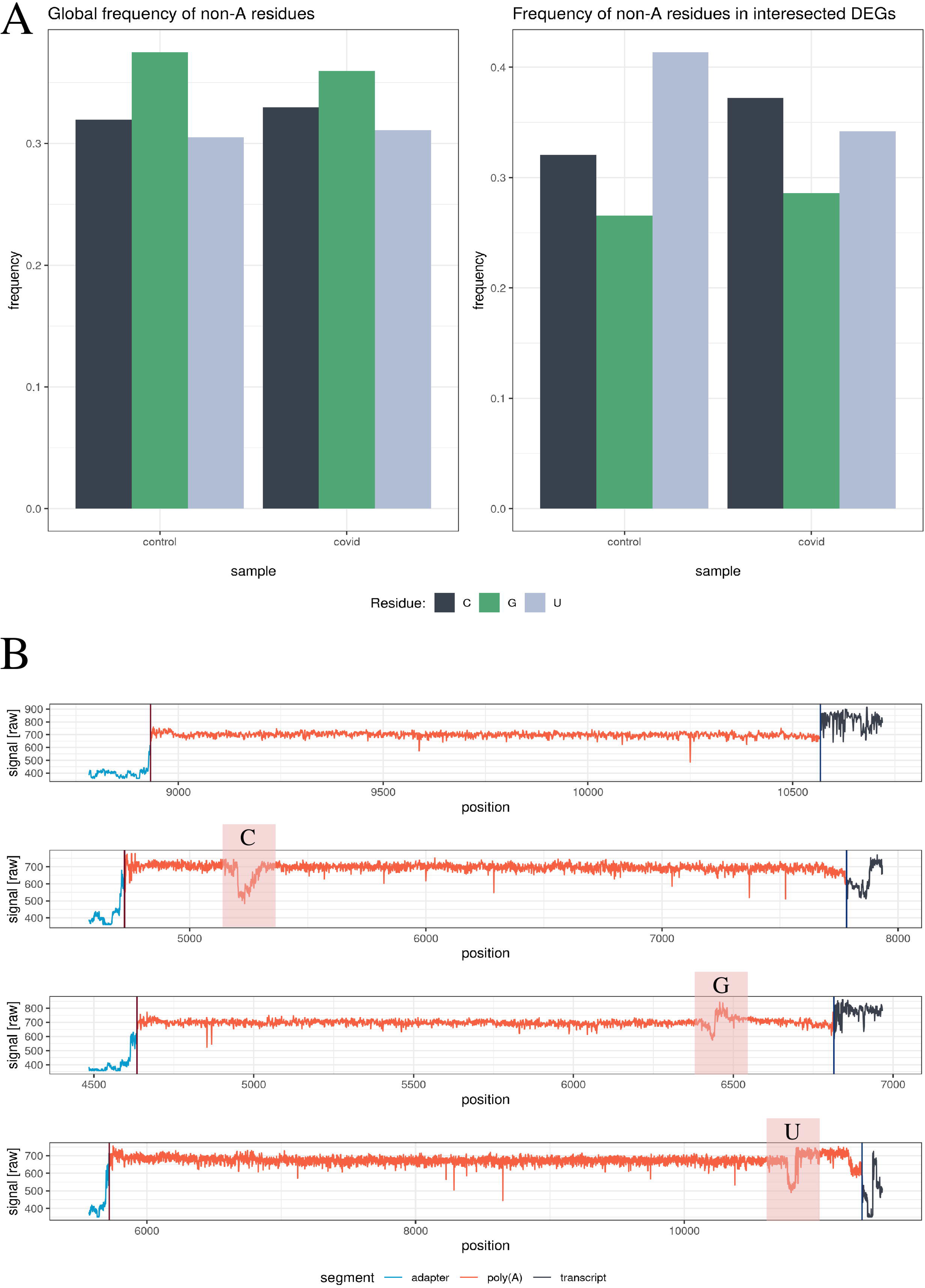
Characterization of Non-A Residues in COVID Samples and Control. **A** Non-A occurrence patterns. The left panel visualizes the overall frequency of non-A residues in control and COVID patient samples. The right panel depicts the distribution of non-A residues within differentially expressed genes (DEGs) shared between both groups. **B** Poly(A) tail illustration. The blue signal represents the adapter, the red signal indicates the poly(A) tail and the black signal corresponds to the transcript. The first panel showcases a canonical poly(A) tail, while subsequent panels demonstrate poly(A) tails with cytosine, guanine, or uracil substitutions, respectively.

### Impact of pseudouridine modifications on RNA in COVID-19

Pseudouridine (psU) was identified as the most common post-transcriptional modification of RNA. A total of 1,214 significant positions were detected in COVID patients and 664 positions in control patients, with 180 positions being common to both groups (Fig. 5A, Supplementary Table 10). These positions were in 1,034 transcripts in COVID patients and 590 transcripts in control patients (Fig. 5B). Positions within genes were also examined, amounting to 788 genes in COVID patients and 469 genes in controls (Fig. 5D). GO analysis of genes with statistically significant psU provided information about the immune response (GO:0006955), response to virus (GO:0009615), and defense response (GO:0006952) as significant processes in the COVID group (Supplementary Table 11). The response to virus (GO:0009615) was not detected as a significant process in the control group, while immune response (GO:0006955) and defense response (GO:0006952) were identified as significant processes (Supplementary Table 12).

**Fig. 5.**
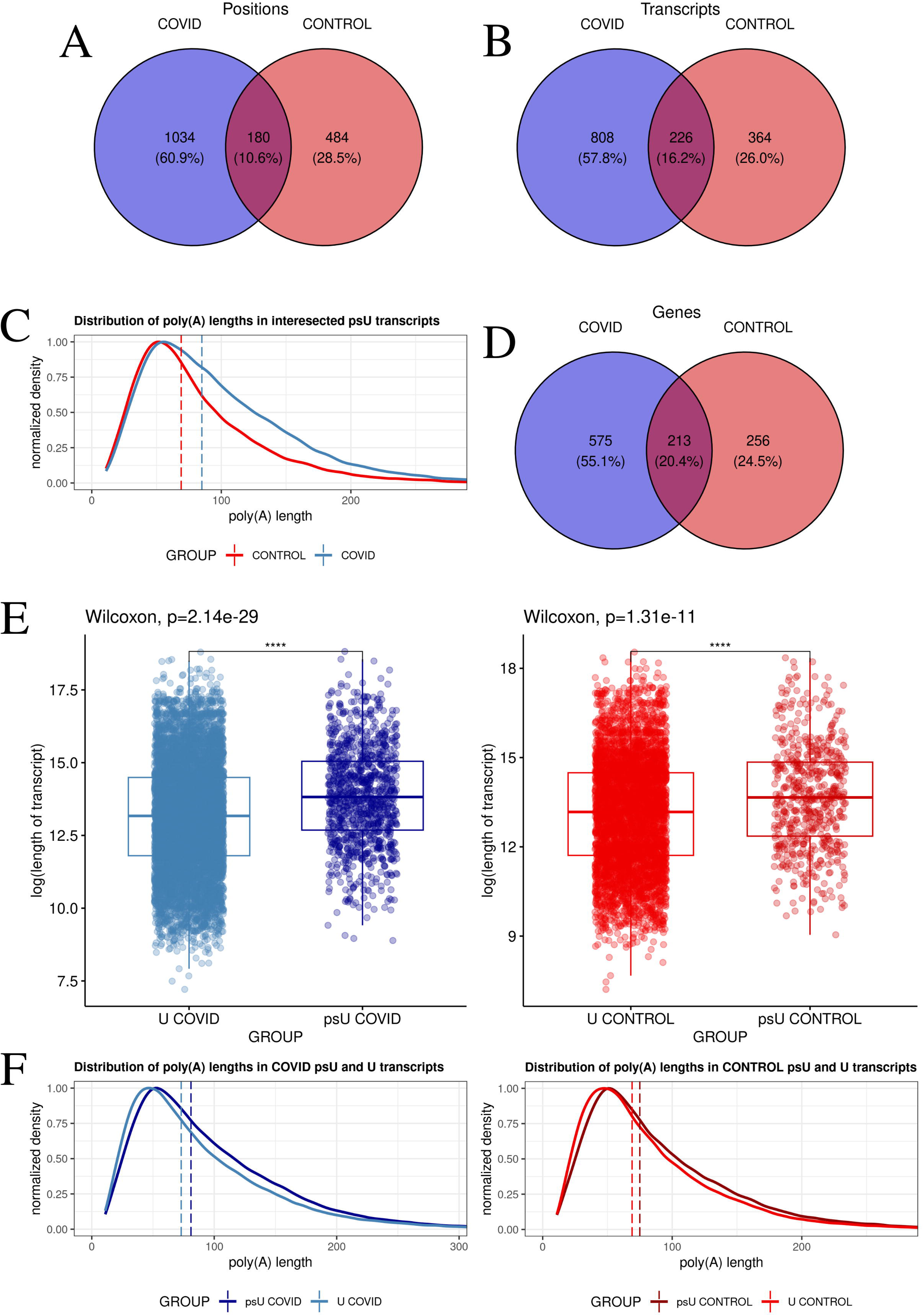
Distribution of pseudouridine and poly(A) tail lengths in control and COVID-19 samples. **A,B,D** Venn diagrams illustrate the occurrence of psU in the control group (red), COVID-19 group (blue), and the overlap between the two groups (dark red) for A positions, B transcripts, and D genes, respectively. **C,F** Density plots depict the normalized transcript density on the y-axis and the length of poly(A) tails on the x-axis. Panel C shows the distribution of poly(A) tail lengths in transcripts with psU present in both groups. Panel F corresponds to the distribution of poly(A) tail lengths in transcripts with psU in the COVID-19 group (left, blue) and in the control group (right, red).

The next step in our analysis involved examining the length of poly(A) tails in transcripts where psU was detected in both groups. A lengthening of poly(A) tails was observed in transcripts with psU in COVID patients compared to control patients (P-value < 2.2e-16) (Fig. 5C). We also decided to examine differences in poly(A) tail lengths between transcripts with psU and uracil in the same genes in both the COVID and control groups. In both cases, a lengthening of the tail was observed in transcripts with psU, with a P-value < 2.2e-16 for both groups (Fig. 5F).

Furthermore, the relationship between the occurrence of psU and transcript length was investigated. Again, transcripts with psU and uracil in the same genes were compared in both the COVID and control groups. Longer transcripts were observed with detected psU compared to those with uracil, with a P-value < 2.2e-16 for the COVID group and P-value = 1.311e-11 for the control group (Fig. 5E, Supplementary Table 13,14).

### Comparative Analysis of m5C Methylation Between Control and COVID-19 Groups

The CHEUI program enabled us to compare m5C methylation sites between the COVID-19 and control groups. A total of 153,845 positions were identified and subjected to differential analysis based on pval_U and stoichiometry_diff. The analysis revealed 689 significant positions within 398 transcripts, which were mapped to 161 genes. 366 positions exhibited stoichiometry_diff < 0.1 (Lower), while 323 exhibited stoichiometry_diff > 0.1 (Higher) (Fig. 6A, Supplementary Table 15). GO analysis revealed significant processes such as the T cell receptor signaling pathway (GO:0050852) and alpha-beta T cell activation (GO:0046631). Molecules with significant m5C did not reveal immune response (GO:0006955), response to virus (GO:0009615), and defense response (GO:0006952) as significant processes (Supplementary Table 16). The genes involved in the observed replacement of hydrogen atoms with methyl groups within RNA molecules were not found to be associated with immunological functions. Unlike pseudouridylation, this modification was not considered to have a direct impact on the organism’s immediate defense mechanisms. Genes *CSF3R*, *CSDE1*, and *KCNAB2* were found to be most enriched with m5C modifications, with 44, 20, and 19 modifications, respectively.

**Fig. 6.**
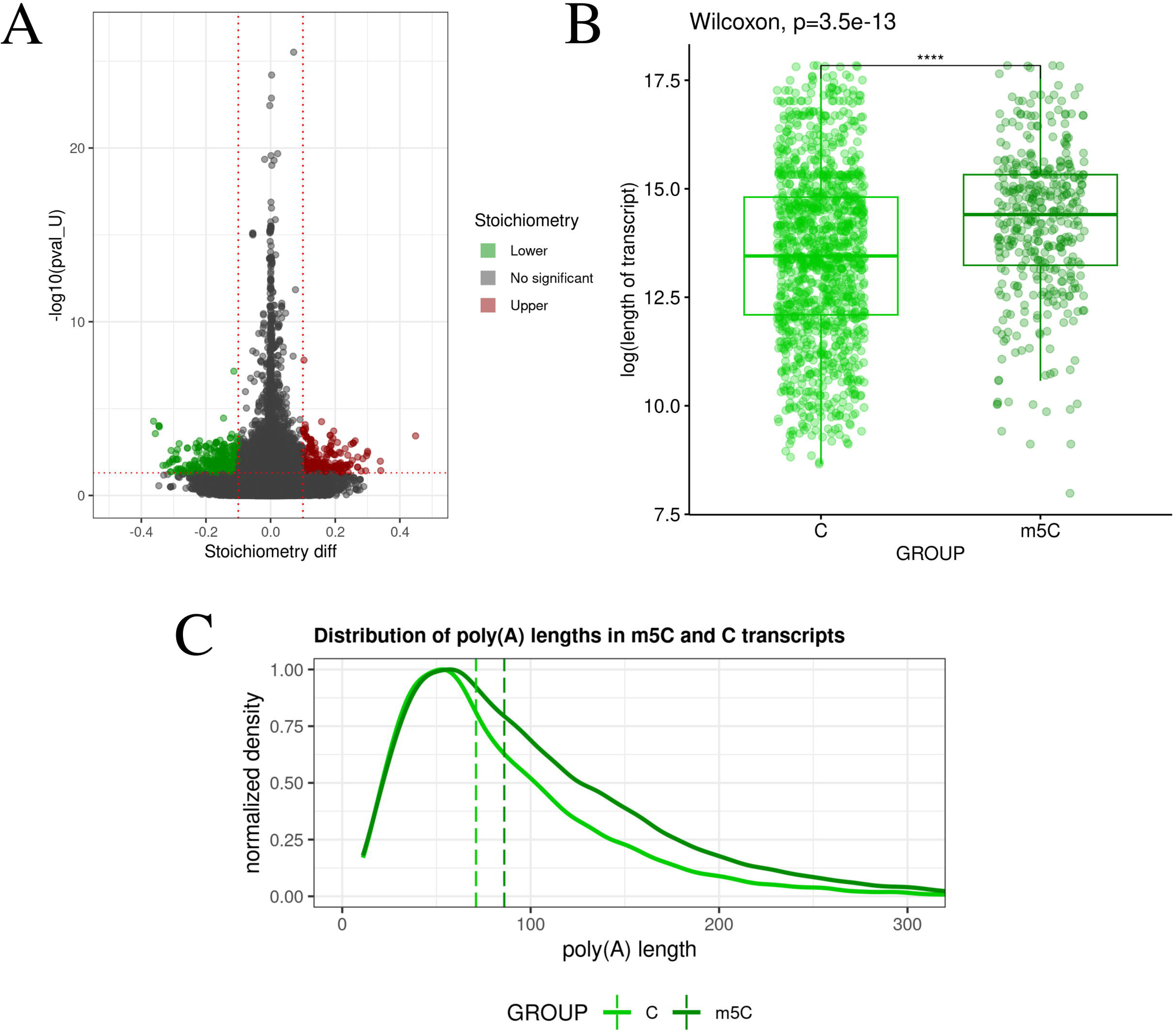
Distribution of m5C methylation and poly(A) tail lengths in control and COVID-19 samples. **A** Volcano plot shows -log10(pval_U) on the x-axis and stoichiometry_diff on the y-axis. Statistically significant positions are colored green (Lower) and red (Upper). Gray dots represent statistically insignificant m5C methylation positions. **B** Boxplots show the relationship between the log2 of transcript length for transcripts with cytosine (light green) and m5C (dark green). **C** Density plot shows the distribution of poly(A) tail lengths in transcripts with cytosine (light green) and m5C (dark green). The y-axis represents the normalized transcript density, and the x-axis represents the length of poly(A) tails.

As a next step in the bioinformatic analyses, the lengths of transcripts and poly(A) tails were examined, similar to the analysis conducted on psU. Transcripts with m5C were observed to be longer than those with cytosine in the same gene (P-value = 3.496e-13), analogous to the findings for psU (Fig. 6B, Supplementary Table 17). Similarly, the lengths of poly(A) tails were found to be longer in transcripts with detected m5C compared to transcripts with cytosine in the same gene (P-value < 2.2e-16) (Fig. 6C, Supplementary Table 18).

### Alternative polyadenylation and its impact on COVID-19

The most frequent alternative polyadenylation (APA) motifs identified were AAUAAA (8 345) and AUUAAA (2 033) (Fig. 7C,F, Supplementary Table 18). APA motifs were found to be most abundant in the three prime untranslated regions (3’UTR) – 11,104 times, in exons - 426 times, in intergenic regions - 402 times, and in introns - 140 times (Fig. 7D). APA sites were observed to occur in various proportions: 1 site was identified in 6 124 genes, 2 sites in 1 613 genes, and 3 sites in 458 genes (Fig. 7E). Analysis of APA site use differences in genes identified 27 significant genes with APA sites, of which 23 were found to have a Δusage value greater than 0.3 and 4 were found to have a Δusage value less than 0.3 (Fig. 7A,B, Supplementary Table 19). Genes exhibiting significantly different APA site usage towards COVID-19 patients included *BMI1*, *JAZF1*, *NIPBL*, and *DDX46*, while in control patients, *RTF1*, *PGS1*, *SPCS3*, and *CCNG2* were identified (Fig. 7A). The GO analysis of genes with significant APA revealed the involvement of six significant processes: (GO:1902494), protein-containing complex (GO:0032991), intracellular protein-containing complex (GO:0140535), nucleoplasm (GO:0005654), nuclear lumen (GO:0031981), intracellular membrane-bounded organelle (GO:0043231) (Supplementary Table 20).

**Fig. 7.**
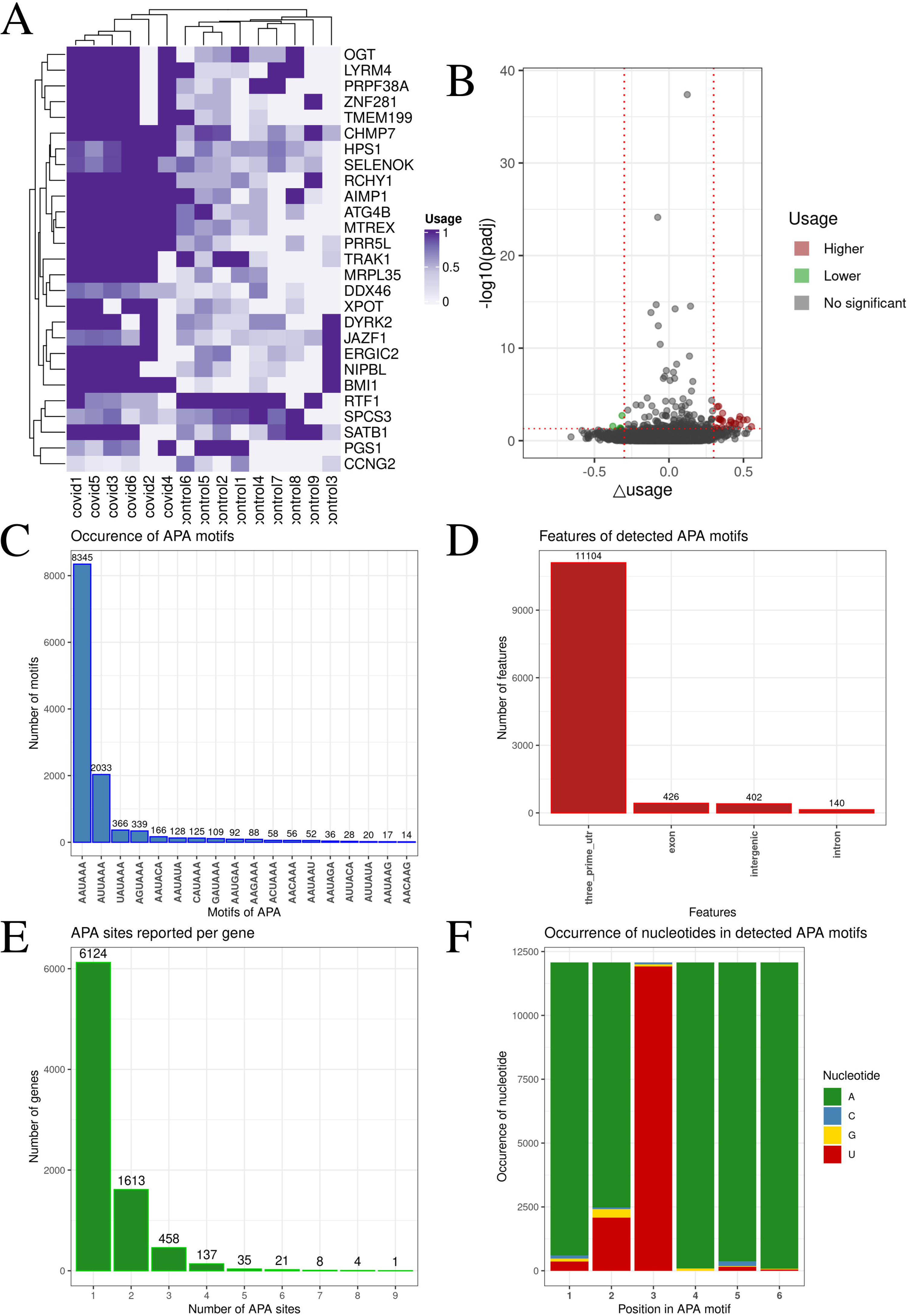
Identification of APA site patterns and associated features. **A** Heatmap visualizes use of APA sites in genes. Darker purple indicates a value closer to 1. Gene names are listed on the right, and patient names are listed at the bottom of the heatmap. **B** Volcano plot shows the relationship between -log10(padj) and Δusage. Gray dots represent statistically insignificant APA sites, green dots represent statistically significant APA sites with Δusage < 0.3, and red dots represent statistically significant APA sites with Δusage > 0.3. **C** Barplots show the frequency of each motif. The x-axis shows the APA motifs and the y-axis shows their count. **D** Barplots show the location of APA sites and their frequency. The x-axis shows the specific location, and the y-axis shows the number of APA sites at that location. **E** Barplots show the number of APA sites per gene. The x-axis shows the number of APA sites, and the y-axis shows the frequency of that number of APA sites in the groups. **F** Barplots show nucleotide frequencies at each position of the detected APA sites. The x-axis shows the position number, and the y-axis the nucleotide frequency. Adenine is represented by green, cytosine by blue, guanine by yellow, and uracil by red.

## Discussion

Differential expression analysis, which involves counting the number of reads per gene in an RNA sequence, has become the primary method for identifying systematic changes across experimental conditions^10^. Studies have shown that combining two sequencing methods can provide additional information through gene expression analysis. Furthermore, it has been shown that these methods could be complementary^11^. The present investigation identified 52 DEGs common to both cDNA and DRS methods. GO analysis of these DEGs categorized them into ‘immune response’, ‘response to virus’, and ‘defense response’ terms. Among the common genes, 2’-5’-Oligoadenylate Synthetase (*OAS1*), Apolipoprotein B MRNA Editing Enzyme Catalytic Subunit 3A (*APOBEC3A*) and Interferon Induced Protein 44 (*IFI44*) were identified. The present study revealed that the SARS-CoV-2 infection was followed by the upregulation of *OAS1* which was earlier recognized as a strong antiviral factor against this virus. It has been found that the inactivation of OAS1’s catalytic activity resulted in the loss of its antiviral function by activating RNase L, which degrades both cellular and viral RNA^12^. Investigations have demonstrated that another gene upregulated in COVID-19 patients, namely *APOBEC3A*, is a critical factor in the inhibition of coronaviruses, by restricting the RNA virus replication^13^. A similar result most probably is exerted by an overexpression of *IFI44* in COVID-19 patients. *IFI44* belongs to interferon (IFN)-stimulated genes and has been previously found to control respiratory syncytial virus (RSV) infection by exerting antiviral and antiproliferative properties^14^.

The primary objective of the present study, however, was not to focus on the differential expression of genes detected in the SARS-CoV-2 infected patients, as we have already described them in our previous study^7^, but to leverage the advantages of RNA long-read sequencing. As Nanopore DRS is based on single-molecule RNA detection, it offers a unique opportunity to examine multiple epitranscriptomic features of individual RNA molecules such as poly(A) tail length, and occurrence of alternative polyadenylation sites, m5C sites and pseudouridine sites, which are often lost during amplification-based RNA-seq techniques and have not been comprehensively explored in COVID-19 patients. Furthermore, the understanding of the functional consequences of RNA modifications has enabled the development of highly effective mRNA-based vaccines for COVID-19^15^. Poly(A) tails were found to be dynamic elements of transcripts, rather than static units simply marking the 3’ end of mRNA, which significantly influence the post-transcriptional regulation of the fragile balance of mRNA survival–degradation^16^. The analysis of these structures may provide novel insights into the post-transcriptional regulation of gene expression in the contexts of development, differentiation, and various disease states^17^. In our study, lengthening of poly(A) tails was observed in as many as 879 genes in COVID-19 patients, compared to only 8 in the control group. The GO analysis of all transcripts with significant poly(A) length changes revealed their significant contribution to terms such as immune response, response to virus and defense response again. This intriguing observation raises a question: why do post-transcriptionally modified transcripts exhibit increased length? As the dynamic part of the poly(A) tail is also longer in these transcripts possibly this elongation may enhance the recruitment of ribosomes and thus facilitate translation initiation. Investigations conducted on *Xenopus laevis* oocytes demonstrated no discernible disparity in the translation of mRNAs with poly(A) tails of 32 or 150 nucleotides in length^18,19^. Moreover, it was found that transcripts with poly(A) tails shorter than 16 nucleotides were not translated. Furthermore, the frequency of molecules possessing fewer than 30 adenines at the 3’ end of the poly(A) tail was significantly lower within the cellular environment. During current research, we identified a change of poly(A) tails median length between the COVID-19 (81 nt) and control patients (62 nt). Moreover, our investigations revealed a significant increase in the lengths of exons, introns, and poly(A) tails within transcripts bearing m5C and psU modifications. The shorter poly(A) tail length observed in modified RNAs suggests a link between the modification and the regulation of the 3’ end^20^. The length of mRNA can influence the density of ribosomes bound to it, which in turn may affect the efficiency of protein synthesis. Shorter gene isoforms are less likely to be targeted by ribosomes compared to longer isoforms, leading to a direct impact on the protein synthesis^21^. Longer genes tend to be associated with functions that are important in the early embryonic stages, while smaller genes tend to play a role in functions that are important throughout the whole life, like the actions of the immune system, which require fast responses^22^. Conversely, extended poly(A) tails may safeguard genes from rapid enzymatic degradation^23,24^. In the present study, both sequencing methods detected significant poly(A) tail elongation in DEGs identified in COVID-19 patients for instance in the Interferon-inducible protein with tetrapeptide repeats 3 (*IFIT3*), Immunoglobulin Lambda Constant 3 (*IGLC3*), Leucine Aminopeptidase 3 (*LAP3*) or Sterile Alpha Motif Domain Containing 9 Like (*SAMD9L*). Genetic or epigenetic variations that promote *IFIT3* expression in response to viral infection may contribute to protection against SARS-CoV-2. IFIT3 restricts viral spread by binding to viral RNA and inhibiting translation^25^. Moreover, IFIT3 amplifies the interferon response by stabilizing the IFIT1 and IFIT2 protein complexes, further enhancing their antiviral effects and upregulating the expression of ISGs, including the previously described OAS1^26–29^. Type I and type II interferons are crucial components of the innate immune response to viral infections, including SARS-CoV-2^30^. A significant upregulation of the IFN-stimulated genes (ISG) has been detected during COVID–19 disease. Overexpression of *IFIT3* has been detected in uninfected or asymptomatic females who were repeatedly exposed to their symptomatic COVID-19 male partners^31^. Given its potential role in antiviral immunity, the *IFIT3* offers a promising avenue for understanding the mechanisms underlying protection from SARS-CoV-2 infection. Elucidating these mechanisms may have significant implications for developing new therapeutic strategies. NSP16 is essential to the SARS-CoV-2 replication cycle because it is essential to coronavirus’ immune evasion^32^. NSP16 is a 2′-O-methyltransferase (2′-O-MTase) that forms part of the replication-transcription complex^33^. Inhibition of NSP16 enhances SARS-CoV-2 susceptibility to IFN-I-induced antiviral effectors, such as IFIT1 and IFIT3^34^. Patients with severe COVID-19 displayed increased B cell activation and upregulation of IGLC3, a marker of antibody processing. These findings suggest a robust antibody response to enhance the host protection and enhanced interferon signaling in these individuals^35^. Single-cell RNA sequencing revealed *IGLC3* among the other 15 differentially expressed genes between patients who survived COVID-19 infection^36^. Accurate identification of prognostic factors in critical COVID-19 patients can aid in risk assessment and guide tailored therapeutic interventions. SARS-CoV-2 infection alters the host gene expression profile, leading to the upregulation of interferon-stimulated genes, including *LAP3*. The interferon IFN-stimulated ISG15 had the largest increase in serum of COVID-19 patients, followed by several other IFN-induced proteins, such as LAP3^37^. Similarly to *SAMD9L* was among the differentially regulated interferon-stimulated genes in mild and severe disease cohorts, suggesting that it may play a critical role in SARS-CoV-2 pathogenesis^38^. Previous studies demonstrated that the SAMD9L pathway acts as a crucial host defense mechanism, which poxviruses actively suppress to establish infection^39^. Notably, this pathway was identified among the interferon-stimulated genes (ISGs) exhibiting significantly reduced expression in patients with severe COVID-19 compared to those with mild cases. The identification of SAMD9L as a downregulated gene in severe COVID-19 highlights its potential role as a critical host restriction factor that SARS-CoV-2 must overcome to establish infection^38^. Moreover in people with severe COVID-19 the infection reveals a diminished antiviral response marked by the downregulation of antiviral genes such as *OAS1*, *SAMD9L* and *IFIT2*, and suppression of antiviral immune response pathways^40^.

Shortening the 3’UTR through alternative polyadenylation (APA) may be a key mechanism contributing to COVID-19 pathogenesis. APA-mediated reduction of 3’UTR length can increase gene expression by evading miRNA-mediated silencing during SARS-CoV-2 infection^41^. Moreover, global 3′UTR shortening affects protein abundance, and the impact of the 3′UTR on protein production may depend on the gene. However, the APA of the genes confers different functions and needs further investigation. It was observed that the expression of 3′ processing factors was down-regulated when cells were infected by vesicular stomatitis virus, which might be one of the reasons underlying genome-wide APA when cells were infected with viruses^41^. It has been found that the expression level of 3′ processing factors is also altered in COVID-19 patients. SARS-CoV-2 proteins can bind to APA factors affecting the gene expression level of APA factors to regulate this process^41^. Additionally, alternative polyadenylation has been suggested to impair antigen presentation by MHC molecules in infected cells. Disrupting alternative polyadenylation and splicing could further enhance the ability of SARS-CoV-2 to evade the host immune response^42^.

The poly(A) tail of mRNA has been conventionally considered a homogenous stretch of adenosine nucleotides, devoid of significant information content beyond its length. However, the non-canonical poly(A) polymerases, TENT4A and TENT4B, have been identified as enzymes capable of incorporating non-A nucleotides, such as guanine, uracil, and cytosine, into the poly(A) tail^43^. While the function of this mechanism remains unclear, it is hypothesized that the presence of these non-adenine nucleotides (non-A mutations) may impede the activity of deadenylase enzymes, slowing the rate of poly(A) tail shortening and increasing the mRNA stability^16,43^ However, Poly(A)-binding protein can stimulate the removal of adenine residues from the poly(A) tail, a process known as deadenylation. This contrasts with the expectation that stable, highly translated mRNAs would possess longer poly(A) tails^23^. It has to be mentioned that the present study revealed also the non-A residues in both COVID and control samples, and SARS-CoV-2-infected patients exhibited increased cytosine content and decreased guanine content in non-A residues.

The methylation of cytosine residue position 5 (m5C), is a widespread RNA modification catalyzed by members of the NOL1/NOP2/SUN domain (NSUN) family (NSUN1-NSUN7) particularly NSUN2 which has the broadest substrate specificity. 5-methylcytosine (m5C) modification was found to play a crucial role in many aspects of RNA processing, tRNA stability, rRNA assembly, and mRNA translation. The effect of viral infection on m5C process modifications is observed. Using bisulfite sequencing, infection with Simian retrovirus was found to affect a total of 2475 m5C sites located on 517 genes related to viral infection^44^. The functional significance of 5-methylcytosine (m5C) modification in viral biology remains largely unknown. SARS-CoV-2 gRNAs carry abundant m5C modifications that affect the virus’s life cycle^45,46^

Low levels of m5C modification in viral RNAs caused by NSUN2 depletion were beneficial to viral replication and infection. Moreover, NSUN2-mediated m5C modification in different regions of viral transcripts reduced the stability and levels of the corresponding mRNA. NSUN2 was found to be a critical host factor in limiting SARS-CoV-2 infection, as NSUN2-deficient mice exhibited more severe infection and lung tissue damage compared to control mice^47^. Certainly, m5C modifications in SARS-CoV-2 transcripts are crucial to viral replication and pathogenicity. In the current investigation, we discovered 689 m5C methylation sites, assigned to the T cell receptor signaling pathway and alpha-beta T cell activation. Jiang et al. ^48^ used methylated RNA immunoprecipitation sequencing to identify m5C modifications in lncRNAs of H1N1-infected and uninfected A549 cells. These authors recognized 2984 differentially modified (DM) m5C sites on lncRNAs in influenza A virus-infected cells, including 1317 upregulated and 1667 downregulated sites, compared to the uninfected group. Moreover, the target genes of methylated lncRNAs were enriched in virus infection-related pathways (e.g. Epstein-Barr virus infection, Measles and Herpes simplex virus 1 infection). In our data, among the analyzed genes, *CSF3R*, *CSDE1*, and *KCNAB2* were found to be most enriched with m5C modifications. *CSF3R* is assumed to be an inflammation-related gene that participates in the immune response to SARS-CoV-2 infection and was significantly upregulated in the blood immune cells of COVID-19 patients^49^. Moreover, it acts as a receptor to promote the activation of granulocytes and macrophages, particularly, which can subsequently upregulate the production of inflammatory cytokines^50^. It has been revealed that CSDE1 is required for translation in human rhinovirus and poliovirus^51^. SARS-CoV-2’s success relies on its ability to repurpose host RNA-binding proteins (RBPs) and to evade antiviral RBPs. Transcriptome analyses identified CSDE1 as a proviral RBP that influences multiple stages of the mRNA lifecycle^52^. CSDE1 knockdown inhibits SARS-CoV-2 replication by suppressing viral RNA levels^52^. Moreover, SARS-CoV-2 may recruit CSDE1 to regulate IRES-dependent translation initiation^53^ of SARS-CoV-2 gRNA and sgRNAs. KCNAB2, a regulatory beta subunit of the potassium voltage-gated channels was recognized as an interferon-stimulated gene upregulated, specific to type I IFN-mediated antiviral signaling upon SARS-CoV-2 infection^54^.

The present study focused also on the pseudouridylation, the modification of uridine to pseudouridine (psU), which is one of the most abundant post-transcriptional modifications of RNA^55^. The current research revealed a total of 1,214 significant psU modification sites in COVID-19 patients and 664 in control patients, with 180 sites shared between the two groups. The GO analysis of genes with statistically significant P-values revealed that the COVID-19 group was significantly enriched for biological processes related to immune response, response to virus and defense response once again. Additionally, a lengthening of poly(A) tails was observed in transcripts with psU in COVID patients compared to control patients. Pseudouridylation has been implicated in regulating both translational processes and cellular stress responses. Alterations in the dynamic pseudouridylation landscape have been previously associated with the pathogenesis of cancer and genetic disorders^56^. The elevated translation rates observed for pseudouridylated mRNA can be attributed to the increased tendency of unmodified mRNA to undergo activation upon binding to RNA-dependent protein kinase^56,57^. Interestingly, present data concerning significant psU modification sites in SARS-CoV-2 infected patients revealed results comparable to these obtained in human HeLa cells^58^. It has been also found, that the strategic incorporation of N1-methyl-pseudouridine (m1Ψ) modifications into mRNA vaccines has been crucial for mitigating their innate immunogenicity and facilitating their efficacy against COVID-19^56,57^. Izadpanah et al.^59^ suggested that SARS-CoV-2 is able to escape the host immune response through the viral RNA pseudouridylation process. According to the current results, the poly(A) tails of pseudouridylated transcripts were statistically longer in COVID-19 (median of 85 nt) compared to the control group (median of 69 nt). We may assume that such a protection mechanism is utilized by viruses to protect against, i.e., rapid degradation of viral transcripts by the host’s enzymes.

## Conclusions

To the best of our knowledge, this is the first study to decipher in such a deep extent both Nanopore long reads and RNA-seq datasets to investigate the whole blood transcriptomic profiles of the SARS-CoV-2 infected patients by providing comprehensive insights into the epitranscriptome features and post-transcriptional modifications. The identification of DEGs such as *OAS1*, *APOBEC3A*, and *IFI44*, as well as genes associated with immune responses, highlights the robust activation of antiviral pathways during COVID-19. Notably, the investigation revealed extensive poly(A) tail lengthening in COVID-19 patients, particularly among immune-related transcripts, suggesting an adaptive mechanism to enhance transcript stability and translation efficiency. Epitranscriptome modifications, including m5C and pseudouridylation, were significantly associated with altered transcript length and immune responses. The identification of specific genes enriched in m5C and psU modifications, such as *CSF3R*, *CSDE1*, and *KCNAB2*, underscores the potential regulatory roles of these modifications in SARS-CoV-2 infection. Exploring further host–SARS-CoV-2 interactions at a deep molecular level may be a fascinating focus for research for future therapeutic treatment.

## Material and Methods

### Patients and sample collection

Peripheral blood samples were collected from nine healthy donors, coded as 1-9, and six patients diagnosed with COVID-19, coded as 10-15. The subjects with confirmed cases of COVID-19 were enrolled at the Clinical Department of Communicable Diseases in Ostróda, Poland. The patients in question ranged in age from 54 to 70 years. They fulfilled the requisite criteria for a viral diagnosis of SARS-CoV-2, with viral genes confirmed by RT-PCR analysis of nasopharyngeal swabs. The RT-PCR reactions were conducted using the commercially available COVID-19 Real-Time Multiplex RT-PCR Kit (Labsystems Diagnostics OY, Vantaa, Finland). The kit is designed to detect the ORF1ab, N, and E genes of the SARS-CoV-2 genome in a single reaction. The RT-PCR reactions were conducted following the manufacturer’s recommended protocol, and the results were analysed using a QuantStudio™ 5 Real-Time PCR System instrument. The inclusion criteria comprised a positive PCR test for SARS-CoV-2, as well as a clinical diagnosis of COVID-19 requiring hospitalisation. The exclusion criteria encompassed patients with neoplasms, autoimmune disease, a pressive or immunodeficient state, and human immunodeficiency virus (HIV) infection. The control group was constituted by volunteers who had tested negative for SARS-CoV-2 infection and showed no signs of respiratory tract infections or lung pathologies, as confirmed by a physician. The control group was constituted in compliance with the following inclusion criteria (verified by a screening questionnaire): absence of a history of travel to high-risk areas, lack of admission to the vaccine, lack of known exposure to a proven or suspected case of SARS-CoV-2 in the previous 14 days, absence of upper or lower respiratory tract infection or any other active illness at the time of blood collection, and lack of past or current history of serious chronic disease such as immune disease. Whole blood samples (3 mL) were collected from all patients (1-15) and placed into Tempus™ Blood RNA Tubes (Applied Biosystems, Waltham, Massachusetts, USA). These samples were stored at -80°C until analysis time. The study was conducted following the ethical standards outlined in the Declaration of Helsinki and received approval from the Bioethics Committee of the Warmia-Mazury Medical Chamber (OIL.3/2021/Bioet) in Olsztyn, Poland. All participants provided written informed consent, as evidenced by their signature, to take part in the study.

### Chest Computed Tomography

Chest computed tomography (CT) was employed as the primary diagnostic tool for managing patients during the initial stages of the severe acute respiratory syndrome coronavirus 2 (SARS-CoV-2) pandemic. A peripheral arterial oxygen saturation level ≤ 93% was one of the criteria used for hospital admission, according to the institutional protocol. All patients underwent a chest CT scan without intravenous contrast in the supine position (Toshiba Medical System, Aquilion Prime type TSX-303A/BK; a tube kilovoltage (kV), 120-135 kV, tube current 530-600 mA, 160 layers, 80 rows). The scans were analyzed using the Osirix MD 11.0™ software (Pixmeo Company, Bernex, Suiça) by two radiologists with experience in chest CT, without previous knowledge of the RT-PCR results of the individual patients. Chest CT scans were qualitatively assessed to identify opacity types, specifying their morphology, distribution, and percentage of involvement of the lung parenchyma.

### Total RNA extraction from peripheral blood

The total RNA was isolated from the whole blood of both the experimental and control groups using the Tempus™ Spin RNA Isolation Kit (Applied Biosystems, Waltham, Massachusetts, USA). Before extraction, the Tempus tubes containing the patient’s blood were thawed and transferred into a 50 mL tube. Subsequently, 3 mL of PBS (Ca²⁺/Mg²⁺-free) was added to reach a total volume of 12 mL. The tubes were vigorously vortexed for a minimum of 30 s and subsequently centrifuged at 4°C at 3000×g for 30 min. Following this, the supernatant was carefully poured off, and the RNA pellet was purified under the manufacturer’s instructions. Finally, total RNA quantity and quality were evaluated utilising an Agilent 2100 Bioanalyzer (Agilent Technologies, USA).

### Nanopore direct RNA sequencing (DRS)

Total RNA isolates were enriched for mRNA using NEBNext® Poly(A) mRNA Magnetic Isolation Module (New England Biolabs) which removed ribosomal RNA (rRNA). Long read libraries were then prepared from 50 ng of poly(A)-tailed mRNA per sample using the Direct RNA Sequencing Kit SQK-RNA002 (Oxford Nanopore Technologies) following the manufacturer’s protocol. SuperScript III Reverse Transcriptase (Thermo Fisher Scientific) was used in the first step of library preparation, which is the synthesis of the complementary strand to the RNA, creating an RNA-cDNA hybrid. Next, sequencing adapters were attached using T4 DNA Ligase (2M U/ml, New England Biolabs) in combination with NEBNext® Quick Ligation Reaction Buffer. The libraries were quantified with the Qubit dsDNA HS Assay Kit (ThermoFisher) and sequenced on a MinION MK1C sequencing device (ONT) using FLO-MIN 106 Flow Cells R.9.4.1 (ONT). The Flow Cells were prepared for sequencing with the Flow Cell Priming Kit EXP-FLP002 (ONT). Long-read digital MinION signals were first converted from POD5 to FAST5 format using the pod5-file-format program (https://github.com/nanoporetech/pod5-file-format). Next, transcriptomic sequences were basecalled by Guppy v.6.0.0 (https://community.nanoporetech.com/docs/prepare/library_prep_protocols/Guppy-protocol/v/gpb_2003_v1_revax_14dec2018/guppy-software-overview).

### Short-read RNA sequencing

RNA sequencing generating short reads was performed by an external service provider, Macrogen (Amsterdam, The Netherlands), utilizing the Illumina technology (Illumina, San Diego, CA, USA). Briefly, the quality (RIN) and quantity of isolated RNA were assessed using the Tapestation 2200 (Agilent Technologies, Santa Clara, CA, USA). Samples with a RIN value greater than 7.0 were selected for further processing. Sequencing libraries were constructed using the Illumina TruSeq Stranded Total RNA with Ribo-Zero Plus rRNA Depletion kit (Illumina, San Diego, CA, USA), following the manufacturer’s protocol outlined in the TruSeq Stranded mRNA Reference Guide (#1000000092426 v01). Generated libraries were quantified using both qPCR and the KAPA Library Quantification Kit (Roche, Pleasanton, CA, USA). Subsequently, libraries were normalized, pooled in equimolar concentrations, and sequenced on the Illumina NovaSeq 6000 platform in a paired-end configuration (2 x 150 bp). Binary base call (BCL) files were converted into FASTQ format using the Illumina bcl2fastq v.2.19 package (https://github.com/brwnj/bcl2fastq). Following conversion, the sequencing data was directed for further analysis.

### Expression profiling based on nanopore DRS

The FASTQ raw reads were then quality-checked and subjected to mapping steps against a reference *Homo sapiens* genome v.GRCh38. This mapping was performed using minimap2 v.2.26 software with *-ax splice* option (10.1093/bioinformatics/bty191). Gene expression profiles were subsequently estimated using featureCounts v.2.0.6 (10.1093/bioinformatics/btt656), based on a GTF file (GRCh38.p14) from which information about hemoglobin coding genes had been removed. A statistical test based on a negative binomial model and shrink, implemented in DESeq2 v.1.42.0^60^ was additionally employed for analysis. The statistical significance of differentially expressed genes (DEGs) was determined using the following parameters: adjusted P-value < 0.05 and |log2(FoldChange)| > 1.

### Expression profiling based on cDNA Illumina sequencing

Raw reads were trimmed using Trimmomatic v.0.39 (https://github.com/usadellab/Trimmomatic)^61^ with the following parameters: *crop: 140, leading: 20, trailing: 20, minlen: 140, avgqual: 20*. Next, STAR v.2.7.11b (https://github.com/alexdobin/STAR)^62^ was used to map FASTQ files against *H. sapiens* reference genome v.GRCh38 utilizing ENCODE standard parameters. Gene count information was obtained using featureCounts with the reference GTF file (without hemoglobin genes). Then, similar to DRS, DESeq2 was used to assess the significance of differential gene expression. The fluctuation of gene expression with adjusted P-value < 0.05 and |log2(Fold Change)| > 1 was considered statistically significant. The Pearson correlation was plotted between the log2(Fold Change) values from cDNA Illumina sequencing and DRS for molecules that showed statistical significance.

### Differential adenylation

The FASTQ files were remapped into *H. sapiens* transcriptome (GRCh38.p14). Tail information for each transcript was then extracted using the nanopolish v.0.14.1 tool (https://github.com/jts/nanopolish). Subsequently, a statistical method based on the Wilcoxon test was applied by the nanotail v.0.1.0 package (https://github.com/smaegol/nanotail) to determine the differences in tail length between molecules transcribed in both conditions. Only reads tagged by nanopolish as ‘pass’ or ‘suffclip’ were considered in the following analyses. mRNA tails shorter than 10 bp and transcripts with fewer than 10 counts were excluded from the analysis. Poly(A) tails of genes with an adjusted P-value less than 0.05 were considered statistically significant. Statistical significance of overall tail length difference between COVID-19 and control patients was assessed using the Wilcoxon test.

### Non-adenine residue analysis

Previously generated nanopolish outcomes, sequencing summary generated by the Guppy basecaller, and FAST5 files were used to identify non-adenine (non-A) sites in the poly(A) tail. This identification was performed by the ninetails v.1.0.0 software (https://github.com/MystPi/ninetails).

### Alternative polyadenylation sites detection

The BAM files generated from mapping long reads to a reference genome were utilized to identify and analyze variations in alternative polyadenylation sites (APA). The LAPA program^63^ was used to predict statistically significant APA sites, which considered the following parameters adjusted P-value < 0.05 and |Δ usage| > 0.3.

### 5-methylcytosine sites detection

The FAST5 files were combined to enable the re-execution of basecalling for both the COVID-19 and control groups. The resulting FASTQ files were re-mapped to the transcriptome sequence using the minimap2 v.2.26 software with the *-ax map-ont* parameter. The resulting BAM files were then sorted using samtools v.1.16.1^64^. Signal data was rescaled to the aligned sequences using nanopolish. The CHEUI tool (https://github.com/comprna/CHEUI)^65^ was used to extract 5-methylcytosine (m5C) differential information. Preprocessing involved predictions of stoichiometry values and modification probabilities at transcriptomic sites using two models. Next, sites that occurred in both control and COVID patients were subjected to m5C differential analysis, in which only sites with P-value < 0.05 and the absolute value of stoichiometry differentiation (stoichiometry diff) > 0.1 were deemed statistically significant.

### Pseudouridine sites detection

The combined FASTQ files from both conditions (COVID-19 and healthy patients) were divided into separate datasets, after which pseudouridine site identification was conducted using the NanoSPA tool (https://github.com/sihaohuanguc/NanoSPA). The reference transcriptome mapped by long reads was analyzed with the following parameters: *remove_intron*, *extract_features*, and *prediction_psU* functions with default options. Pseudouridine sites with a probability > 0.95 were considered statistically significant.

### Functional annotations

All statistically significant molecules were then scanned for enrichment analysis in Gene Ontology (GO) annotations^66,67^ using the g:profiler v.0.2.2 R package^68^. For the essential genes, biological processes (BP), cellular components (CC) and molecular functions (MF) terms were assigned as ontological annotations. Enrichment analysis was subsequently employed to identify GO terms regulated by significant molecules, using an adjusted P-value < 0.05 cut-off.

### Transcript length fluctuation

Transcript length information was obtained from the reference GTF (*H. sapiens* v.GRCh38). Differences in transcript lengths between transcripts with detected significant RNA modifications and the other transcripts of the same gene were compared using the Wilcoxon test. A significance level of P-value < 0.05 was set.

### Visualization

The visualizations were generated using the R environment and the following packages: ggplot2 v.3.5.1^69^, ComplexHeatmap v.2.18.0^70^, and ggvenn v.0.1.10 (https://github.com/yanlinlin82/ggvenn). Furthermore, visualizations offered by the previously employed software were leveraged.

### Data availability

The data underlying this article are available in the European Nucleotide Archive repository, under accession numbers PRJEB84380, PRJEB74103 (https://www.ebi.ac.uk/ena/browser/view/PRJEB84380, https://www.ebi.ac.uk/ena/browser/view/PRJEB74103).

## Contributions

Conceptualization, M.A.M., and M.M.; Formal Analysis, M.A.M.; Software, M.A.M., E.L., K.G.M., and M.M.; Methodology, M.A.M., K.K., J.S., and M.M.; Investigation, M.A.M., K.K., E.L., Ł.P., K.G.M., P.I., J.S., L.G., and M.M.; Resources, B.M., P.I., P.K., L.G., and M.M.; Writing - original draft: M.A.M., K.K., E.L., K.G.M., P.I., J.S., L.G., and M.M.; Supervision M.M.; Project administration M.M.; Funding acquisition J.S. and L.G.; All authors read and approved this manuscript.

